# New therapeutic combination to enhance endocytosis of antibodies and nucleic-acid aptamers targeting EGFR in glioblastoma cells

**DOI:** 10.1101/2024.10.22.617611

**Authors:** Elisabete Cruz Da Silva, Charlène D’Ancona, Hélène Justiniano, Valérie Calco, Dilara Sensoy, Pascal Villa, Romain Vauchelles, Maxime Lehmann, Laurence Choulier

## Abstract

Active targeting is based on the binding of ligands to receptors present on the surface of targeted cells, in order to promote the internalization of the drugs conjugated to the ligands. Several conjugates are already in use or under development for active targeting of tumors, the most widely known being antibody-drug conjugates (ADC). They combine the specificity of monoclonal antibodies with the cytotoxicity of chemotherapeutic molecules. Other than antibodies, nucleic-acid aptamers, are promising ligands to deliver conjugated drugs by active targeting in tumor cells. The therapeutic efficacy of conjugates largely depends on their endocytosis and vesicular trafficking. However, so far, no therapeutic approach to enhance endocytosis of conjugates is available. In recent studies, we showed that gefitinib, a tyrosine kinase inhibitor directed against the epidermal growth factor receptor EGFR, induces a massive, non-physiological endocytosis of EGFR, known as gefitinib-mediated endocytosis (GME), in different glioblastoma cell lines. We thus hypothesized that besides promoting endocytosis of EGFR, gefitinib could also promote endocytosis of its ligands. In this study, we proved by quantitative fluorescence bioimaging, that gefitinib is indeed able to strengthen the endocytosis of fluorophore-conjugated EGFR-specific antibodies and aptamers. We also showed that the GME potentiates the toxicity of an antibody-drug conjugate, even at low concentrations. Our results suggest the development of a new therapeutic combination, of ADC and gefitinib, to potentiate the delivery of ADC and likely other conjugates targeting EGFR in glioblastoma, while limiting side effects on non-targeted cells.

## 1. INTRODUCTION

Glioblastoma (GBM) is the most aggressive form of brain tumor. While progress on understanding the biological mechanisms of GBM is reported (Fares et al., 2024), treatment strategies have remained largely unchanged since 2005 (Stupp et al., 2005). Extremely invasive, GBM is particularly resistant to surgical resection and to current approaches to radiotherapy, chemotherapy and targeted therapy. The epidermal growth factor receptor (EGFR/HER1/Erb1) is a tyrosine kinase receptor, playing a pivotal role in driving the tumor’s progression and aggressiveness (Westphal et al., 2017). Amplification of the HER1 gene (EGFR/ErbB1) is detected in 40 to 55% of GBMs (Furnari et al., 2015). Therapies targeting EGFR currently being used and improving patient care, in other cancers like metastatic colorectal cancer, head and neck squamous cell carcinoma, through antagonistic monoclonal antibodies (e.g., Cetuximab, ERBITUX) or non-small cell lung cancer through EGFR tyrosine kinase inhibitors (e.g., gefitinib, IRESSA), have proven ineffective in clinical trials for GBM patients (Cruz Da Silva et al., 2021b), while the causes of resistance to treatment have not yet been identified, highlighting the need for further strategies to develop therapeutic approaches (Pan and Magge, 2020).

While the signaling properties of EGFR and the signaling cascades lose attraction for GBM therapy, EGFR becomes attractive as a docking molecule. Its internalization and intracellular trafficking are critical events in cell signaling and regulation enabling to carry effector agents (Westphal et al., 2017). Upon binding to its ligands, such as the epidermal growth factor EGF, EGFR dimerizes and auto-phosphorylates, triggering endocytosis. The internalized receptor-ligand complex is then transported to early endosomes, where it can either be recycled back to the plasma membrane or degraded in lysosomes (Caldieri et al., 2018). The balance between recycling and degradation of EGFR is critical for receptor signaling, and dysregulation of EGFR trafficking is associated with various pathologies, including cancer. Altering the expression of proteins that regulate EGFR trafficking leads to continuous downstream signalling (Caldieri et al., 2018). In GBM, EGFR endocytosis and membrane trafficking, key players in EGFR signaling process (Sousa et al., 2012), are often deregulated (Colin et al., 2019; Grandal et al., 2007; Kondapalli et al., 2015; Liu et al., 2021; Park et al., 2018; Yang et al., 2019; Ying et al., 2010), highlighting the importance of these processes in tumour progression and aggressiveness (Chang et al., 2013; Kondapalli et al., 2015; Walsh et al., 2015, p. 2; Walsh and Lazzara, 2013; Ying et al., 2010; Zhou et al., 2017). Indeed, the dynamic process of endocytosis and intracellular trafficking has gained significant progress these last years with the development of specific ligands for the treatment of cancer cells, including antibodies (Kalim et al., 2017), but other ligands may also hold therapeutic promise. In this manuscript, we will focus on antibodies and nucleic acid aptamers targeting EGFR.

In the field of antibody-based targeted therapies, antibody-drug conjugates (ADCs) directed against EGFR opens up new therapeutic perspectives for the treatment of GBM (Hui K. Gan et al., 2023). ADCs combine the selectivity of a monoclonal antibody with the toxicity of conjugated cytotoxic molecule by a linker, enabling its targeted delivery. ADCs are administered intravenously. They recognize antigens highly expressed at the surface of the targeted cells with limited heterogeneity across the tumour and with low normal tissue expression to limit collateral cytotoxic effects, and should be internalized by receptor-mediated endocytosis (RME) (Diamantis and Banerji, 2016). In the lysosomes, the release of the cytotoxic payload via cleavage of the linker or degradation of the ADC, kills or halts the proliferation of the targeted cell (Kalim et al., 2017). In recent years, improvements in chemical techniques have enabled the combination of extremely powerful cytotoxic agents (immunotoxins, radioisotopes, cytotoxic agents against intracellular targets like DNA and the tumour cytoskeleton (Hui K Gan et al., 2023)), leading to the development of more effective and safer ADCs, and resulting in highly selective and cytotoxic anti-cancer treatment. However, ADC clinical efficacy is hindered by several factors, including lower internalization efficiency (Takahashi et al., 2022). Enhancing internalization efficiency is crucial to improve ADCs therapeutic efficiency (DeVay et al., 2017; Kalim et al., 2017).

Aptamers are oligonucleotides referred as chemical antibodies (Zhou et al., 2016), binding to their targets with high affinity and specificity. Despite their clinical potential, illustrated by ongoing trials, of which two in oncology, and two FDA approved for wet, age-related macular degeneration (AMD) and geographic atrophy secondary to AMD (Mullard, 2023), the therapeutic potential of aptamers for targeted therapies is still in its infancy (Mahmoudian et al., 2024). For active targeting, aptamers provide unique advantages over antibodies, including their smaller size (10-30 kDa) which allows for deeper penetration into tumours (Xiang et al., 2015), their lower immunogenicity, lower production cost, and easier synthesis and modification. This last point is particularly interesting as coupling to different kind of cargoes, is rendered extremely precise, reliable, and reproducible. In addition to delivering small molecules (e.g. drugs, toxins), the versatility of aptamers enables precise delivery of other cargoes like nucleic acids, proteins, peptides, nanoparticles. Aptamers targeting cell-surface receptors internalized by RME are innovative and ideal tools for studying intracellular trafficking in tumor cells.

In different GBM cell lines, we have recently shown, that a first-generation EGFR tyrosine kinase inhibitor (TKI), gefitinib (Singh et al., 2023), approved by the FDA in 2003, strongly increases EGFR endocytosis and associated proteins (Blandin et al., 2021; Cruz Da Silva et al., 2021a). This process, called gefitinib mediated endocytosis (GME), induces EGFR accumulation in early endosomes, involving canonical endocytosis proteins (dynamin 2, Rab5) (Blandin et al., 2021). Based on these results, and since the efficacy of EGFR targeted therapies depends on intracellular delivery, we investigated how and if GME would enable endocytosis of EGFR specific ligands, potentially enhancing the efficacy of drugs conjugated to these ligands. In our study, we mainly focused on the EGFR-targeted antibody cetuximab and nucleic acid aptamer E07 (Li et al., 2011). Cetuximab, approved by the FDA in 2004, is used in the clinical management of metastatic colorectal cancer and advanced squamous cell carcinoma but has not yet been employed as a tumor-targeting antibody in anti-GBM ADCs. We recently demonstrated that the E07 aptamer, which is particularly suited for cargo delivery in various EGFR-positive cells (pancreatic (Ray et al., 2012) and epidermoid carcinoma (Li et al., 2011)), binds and is internalized in EGFR-positive GBM cells, but not in cells that do not express EGFR (Cruz Da Silva et al., 2022).

Using quantitative fluorescence microscopy, our results revealed that gefitinib similarly promotes the endocytosis of EGFR ligands. Building on these findings that combining cetuximab and gefitinib selectively enhanced cetuximab uptake by EGFR-positive GBM cells, we used a clinically approved cetuximab-based ADC to validate this drug-combination. Our results demonstrated a cellular cytotoxic synergy between the ADC and gefitinib, indicating that this dual therapy while enhancing the administration of ADC, might enable effective doses to be reduced, to the benefit of patients. Altogether, our results highlight a novel mechanism whereby GME promotes the uptake of different EGFR-targeting nucleic acid aptamers, antibodies and ADCs, demonstrating that combination with gefitinib can potentiate the delivery and efficacy of different EGFR-targeted ligands. Indeed, they provide strong support for ongoing clinical approaches where combining therapies are opening new perspectives (Cruz Da Silva et al., 2021b; Menon et al., 2022; Shikalov et al., 2024).

## 2. MATERIALS AND METHODS

### 2.1. Materials

Gefitinib (ZD1839) was obtained from Med Chem Express (Monmouth Junction, USA) and Cayman Chemical (Ann Arbor, USA). Nucleic-acid aptamers were purchased from IBA Lifesciences (Goettingen, Germany) and Eurogentec (Seraing, Belgium). The sequence of the 2’-fluoro-modified pyrimidine 48mer RNA aptamer E07 (Cruz Da Silva et al., 2022; Kratschmer and Levy, 2018; Li et al., 2011) targeting EGFR, conjugated to cyanine 5 in 5’ (Eurogentec, Belgium) has been described elsewhere (Cruz Da Silva et al., 2022). The mouse antibody AY13 conjugated to PepCP-cyanine5.5 was obtained from BioLegend (San Diego, USA). Cetuximab was conjugated to AlexaFluor 647 according to the manufacturer’s instructions (A20186, Thermo Fisher Scientific, Dardilly, France). The antibody-drug conjugate, composed of cetuximab-APN-PEG_5_-VC-MMAE (with APN for arylpropiolonitrile, PEG for polyethylene glycol, VC for valine-citrulline, MMAE for monomethyl auristatin E), with drug-to-antibody-ratio of 4.3 determined by hydrophobic interaction chromatography, was synthetized by Syndivia (Strasbourg, France).

### 2.2. Cell culture

Cell culture media and reagents were from Lonza (Basel, Switzerland) and Gibco (Thermo Fisher Scientific, Waltham, MA, USA). The human GBM cell lines U87MG EGFR WT (named U87 in the manuscript), and LN319 were kindly provided by Dr. Frank Furnari (Inda et al., 2010; Nishikawa et al., 1994) and Pr. Monika Hegi (Lausanne, Switzerland), respectively. The cell lines were maintained in a 37 °C humidified incubator with 5% CO2 in Eagle’s minimum essential medium (EMEM) with 10% fetal bovine serum (FBS). The culture medium for U87 was supplemented with 1% sodium pyruvate, and 1% non-essential amino acids.

### 2.3. Confocal microscopy analysis of fluorescent ligand internalization and colocalisation

Adherent cells were plated on sterile glass coverslips, for one night at 37°C in culture medium, washed twice with PBS-cation buffer (phosphate-buffered saline, 1 mM MgCl_2_, 0.5 mM CaCl_2_; pH 7.4), and then incubated for 1 h in OptiMEM (Gibco) or PBS-cation buffer containing 3% bovine serum albumin (BSA). For experiments involving aptamers, labeled aptamers were denatured at 95 °C for 3 min, incubated on ice for 5 min before being resuspended at 100 nM final concentration in PBS-cation buffer. For experiments involving all ligands, fluorescent antibodies at 10 µg/ml (# 66 nM) or aptamers at 100 nM were applied to cells for 1 h at 4 °C in humid chamber in the dark. Cells were then washed in OptiMEM or PBS-cations, treated with gefitinib at defined concentrations (0, 0.5, 5, 10 or 20 μM) for specified period of time (from 0 to 24h) at 37°C. For the negative controls, gefitinib was substituted with Dimethyl Sulfoxide (DMSO). Coverslips were then washed and remaining cell-surface ligands were removed by acid wash (sodium acetate 0.2M, pH 2.8 for 5 min on ice) (Blandin et al., 2021). After washing, cells were fixed for 5 min in 4% paraformaldehyde (PFA), permeabilized for 2 min with 0.1 % Triton, washed again and incubated in PBS containing BSA 3% for 30 minutes. Then, immunocytochemistry was performed with the following primary antibodies: rabbit anti-Rab5 to label early endosomes (clone C8B1; Cell signalling technology; 1/1000) and mouse anti-CD63 to label late endosomes (clone H5C6, DSHB, 1/50). Primary antibodies were added overnight (O/N) at 4 °C, followed by washes and incubation for 1 h at RT with a secondary antibody conjugated to Alexa 488 or 568 (anti-rabbit Alexa 568, A11011, Invitrogen, 1/2000 and anti-mouse Alexa 488, A11029, Invitrogen, 1/2000). 4’,6-diamidino-2-phenylindole (DAPI) was added at 1 µg/mL for 45 min to visualize nuclei. Washing steps were performed before mounting using fluorescent mounting medium (S3023; Dako, Carpinteria, CA, USA).

### 2.4. Image analysis

Images of apta- and immunofluorescence were acquired using a Leica TCS SPE II confocal microscope at 63× (oil immersion) magnification. Experiments were performed as technical replicates with a minimum of 3 independent repeats. For each experiment, identical background subtraction and scaling were applied to all images. Microscopy images were processed using ImageJ 1.53t software with 2–4 cells per image with an average of 50 cells / condition. The contour of each cell was manually delimited and the mean integrated fluorescence intensity was measured using ImageJ software as previously described (Blandin et al., 2021; Fechter et al., 2019). Pearson correlation coefficients were determined using JACoP plugin ImageJ software (Bolte and Cordelières, 2006; Dunn et al., 2011).

### 2.5. Cell viability assays

Cells were seeded at 1.5×10^4^ cells/well in 96-well plates (Nunc Edge 2.0, ref 15533115, Thermo Fisher Scientific). After 24h of incubation (cell incubator, 37 °C, 5% CO2), cells were washed and treated in triplicate by addition of 50 µl of medium containing the following drugs: gefitinib à 2 and 20 µM, MMAE 1µM, MMAE 1µM + gefitinib 2 µM, MMAE 1µM + gefitinib 20 µM. An internal control of cytotoxicity with 50 µM chlorpromazine (Sigma C8138) was included in all experiments (Calas et al., 2020; Dudani and Gupta, 1987). The dose-response curves were built using the following final concentrations: 10^−10^, 10^−9^, 10^−8^ and 10^−7^ M of cetuximab and ADC, alone or with Gefitinib treatment at 2 µM and 20 µM. For controls, DMSO concentration (1% final) was the same in all wells. All treatments were done using a fully automated robotized platform (Beckman Coulter, Villepinte, France). Cell viability was measured after 72h of treatment using the WST-1 assay (WST-1 Cell Proliferation Assay System, ref MK400, TaKaRa) according to the supplier’s protocol. Briefly, WST-1-containing medium was added to cells and cell viability was determined by measuring absorbance at 450 nm using an Envision reader (Perkin Elmer, Villebon-sur-Yvette, France) 2h after incubation at 37°C and 5% CO_2_. Each measurement was performed from four independent experiments and results were expressed as means of independent experiments.

### 2.6. Statistics

Bioimaging data were analyzed with GraphPad Prism version 5.04 and are represented as means ± SEMs. The statistical analysis of data was performed with One-way ANOVA with Bonferroni correction, whereas the student t-test was used for direct comparison between two conditions. Significance level is controlled by a 95% confidence interval. Results are considered significantly different when p < 0.05. **p<0.01, ***p<0.001, **** p<0.0001. For cytotoxic assays, statistical calculation of Z’ (Zhang et al., 1999) was performed as previously described (Calas et al., 2020). In our experiments Z’ was greater than 0.5, which is the proof of robustness.

## 3. RESULTS

### 3.1. Gefitinib enhances the internalization of anti-EGFR antibodies in a dose-dependent manner

In recent studies, we showed that tyrosine kinase inhibitors, including gefitinib, induced EGFR accumulation in early-endosomes of glioma cells as a result of an increased endocytosis, and increased the internalization of one of its natural ligand EGF (Blandin et al., 2021; Cruz Da Silva et al., 2021a). We thus hypothesized that gefitinib-based EGFR internalization might as well promote the internalization of other ligands.

We investigated this effect on three GBM cell lines, U87MG, T98 and LN443. After ensuring by immunoblot that these three cell lines express EGFR and that EGFR expression is not altered after 20 µM gefitinib treatment upon time (from 0 to 24h, Figure S1), we studied the effect of gefitinib on the internalization of the commercially available anti-EGFR antibody AY13 conjugated to the fluorescent probe Cyanine5 (AY13-Cy5). The three cell lines were incubated at 4°C in the presence of mAb AY13-Cy5 at 10 µg/ml, then treated with different doses of gefitinib (0, 0.5, 5, 10 and 20 µM) at 37°C for 4 h. After acid washing to remove surface antibodies, the cells were fixed and confocal microscopy was used to observe the inner presence of antibodies (Figure 1A). Without gefitinib, intracellular antibody labelling is weak. Remarkably, after 4 h of treatment, gefitinib provoked a massive intracellular internalization of the anti-EGFR antibody, indicated by the intracellular punctuated structures. Indeed, these structures were enlarged compared to cells without gefitinib treatment, likely due to excess membrane fusion, as already described for gefitinib-mediated EGFR internalization (Blandin et al., 2021). Quantification of antibody internalization revealed that with increasing concentrations of gefitinib, progressive antibody internalization was observed, with a similar profile in all three cell lines. At 20 µM gefitinib, the fluorescence intensity reflecting AY13-Cy5 internalization increased by 7-, 18- and 21-fold in U87, LN443 and T98 cells, respectively, compared with untreated cells (Figure 1B). Indeed, gefitinib, by enhancing EGFR endocytosis, also strongly increase internalization of this anti-EGFR antibody.

**Figure 1.**
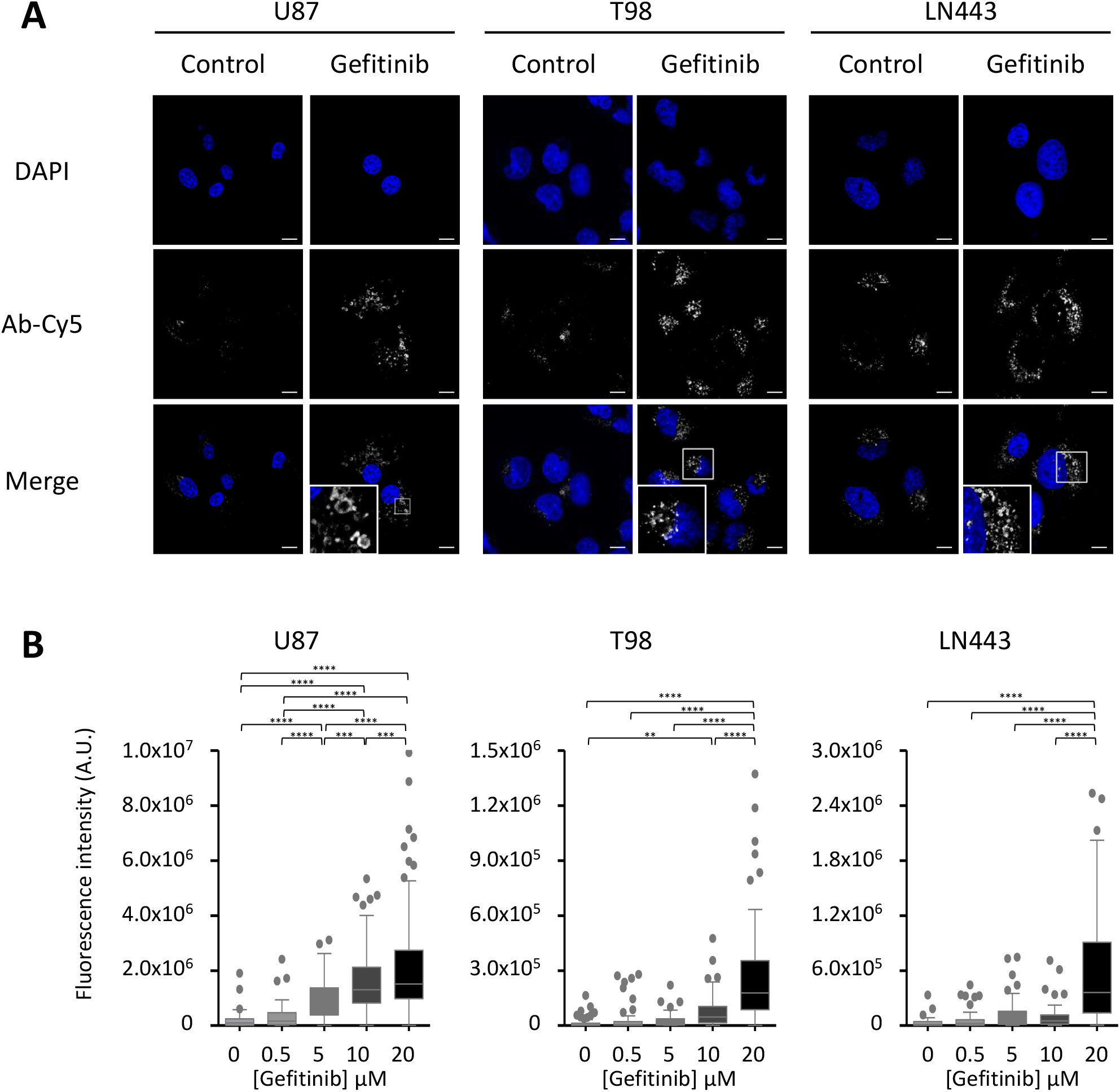
Concentration-dependent enhancement of anti-EGFR antibody internalization by gefitinib in U87, T98 and LN443 GBM cell lines. Cells were incubated at 4°C in the presence of the anti-EGFR mAb clone AY13 conjugated to the fluorophore cyanine 5. They were then incubated with gefitinib at different concentrations (0, 0.5, 5, 10 and 20 µM - 0 corresponds to DMSO treatment) in culture medium, at 37°C for 4 hours. **A**. Representative images. The responses from the cyanine5-conjugated antibody (Ab-Cy5) and DAPI are shown in white and blue, respectively. Scale = 12 µm. High-magnification images are from inserts in the perinuclear zone. **B**. Quantification of anti-EGFR AY13-Cy5 antibody internalization. Fluorescence intensity is expressed in arbitrary units and represented as Tukey boxes using Graphpad Prism software.

As the internalization enhancement happened on the three EGFR-expressing cell lines, we further studied the effect on the U87 cell line with gefitinib at the concentration of 20 µM, in accordance with the results shown in Figure 1 and with our previous studies (Blandin et al., 2021). As an alternative to the mouse antibody AY13, we focused on cetuximab, an antibody targeting EGFR widely used in clinic and highly relevant to human health.

### 3.2. Gefitinib localizes cetuximab in endosomes of EGFR-positive glioblastoma cells

We first explored whether gefitinib, like its effect on the internalization of the antibody AY13-Cy5, also promotes the internalization of cetuximab into U87 GBM cells. Bioimaging experiments akin to those in Figure 1 were conducted, using cetuximab-conjugated to Cyanine 5 (Cetuximab-Cy5), and expanding our study by testing different time points: 15 minutes, 1 hour, 3 hours, and 24 hours. We compared the EGFR-positive cell line U87 with the EGFR-negative LN319 cell line. These two cell lines were selected due to their well-characterized EGFR expression levels, which our research team determined in a recent study (Cruz Da Silva et al., 2022). This characterization involved western blotting, confocal microscopy, and flow cytometry with various anti-EGFR antibodies, including Cetuximab-Cy5.

Representative confocal images of fluorescent Cetuximab-Cy5 and fluorescence quantifications are shown in Figures 2A-B. In EGFR-negative LN319 cells, irrespective of the treatment time, the absence of visible fluorescence intensity from Cetuximab-Cy5 indicates that the low level observed in quantitative experiments corresponds to fluorescence noise. However, in the EGFR-expressing U87 cell line, cells display the presence of Cetuximab-Cy5. In absence of gefitinib, the fluorescence intensity rises smoothly over time, to 1.7x after 24h compared to 15 minutes, indicating that Cetuximab is progressively internalized inside the EGFR-positive cell line. Under gefitinib treatment, the fluorescence intensity strongly increases, to 1.4x and 1.7x at 3h and 24h respectively, compared to untreated U87 cells at the same time points.

**Figure 2.**
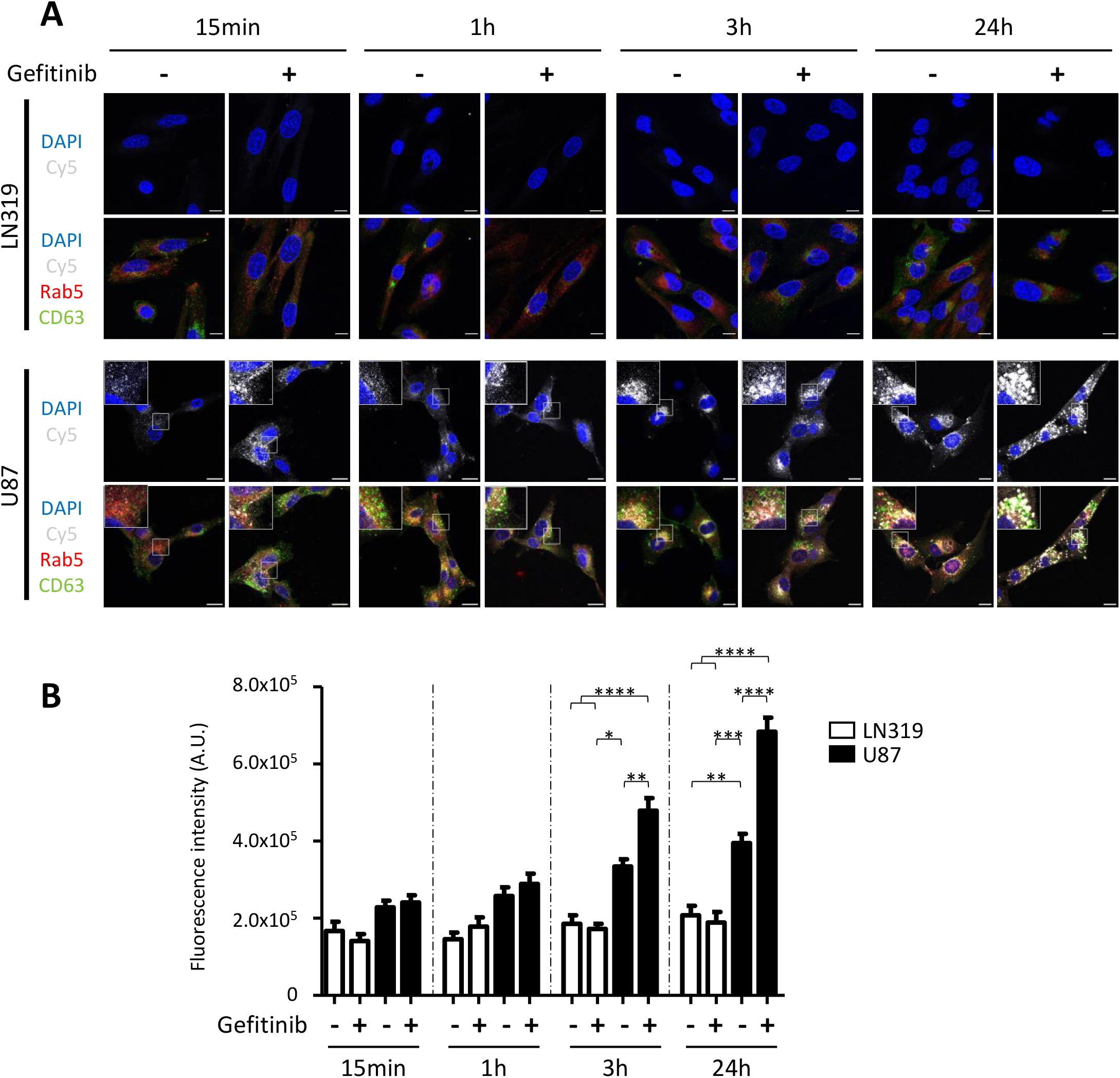
Effect of gefitinib treatment on the internalization and co-localisation of the cyanine5-conjugated cetuximab in GBM cell lines. LN319 and U87 GBM cell lines were incubated for 1h at 4°C in the presence of 10 µg/ml of cyanine5-conjugated cetuximab (Cetuximab-Cy5), and then with 20 µM gefitinib or DMSO (control) in culture medium, at 37°C for different times (15 min, 1h, 3h, 24h). Cells are then acid washed, fixed, immunolabeled and the nucleus are stained with DAPI. **A**. Representative confocal images, showing DAPI (blue to label nucleus), Rab5 (red to label early endosomes), CD63 (green to label late endosomes), and Cetuximab-Cy5 (grey). Magnified images of U87 cells are from the inserts. Scale bar = 12 μm. **B**. Quantification of the Cetuximab-Cy5 internalization. Fluorescence intensity is expressed in arbitrary units.

To determine the intracellular location of Cetuximab-Cy5, immunostaining experiments were performed, using the vesicular trafficking proteins Rab5 and CD63. Similar to its internalization in HeLa cells (Takahashi et al., 2022), endocytosed cetuximab colocalized with early endosome biomarkers, from as early as 15 min. As shown in Figure 2A, gefitinib triggered massive cetuximab localization close to the cell nuclei, in large Rab5- and CD63-positive early and late endosomes. Colocalization quantification indicates that the intracellular localization of Cetuximab is similar regardless of the presence or absence of gefitinib treatment in early and late endosomes (Figure S2). Altogether, these data highlight that, in cells that express EGFR, gefitinib substantially and rapidly enhances intracellular internalization of cetuximab, condensing cetuximab inside large endosomal intracellular compartments.

### 3.3. Gefitinib enhances nucleic acid aptamer internalization and localizes aptamers in endosomes of EGFR-positive glioblastoma cells

We examined whether gefitinib enhance the internalization of other ligands than antibodies. We chose nucleic acid aptamers due to their potential as promising tools for targeted delivery (Mahmoudian et al., 2024) and our team’s expertise in the identification and characterization of aptamers targeting cell-surface receptors (Cruz Da Silva et al., 2022; Fechter et al., 2019; Mercier et al., 2017). We realized the same experiments than those described with the Cetuximab-Cy5, using the EGFR-specific aptamer E07, conjugated to the Cyanine 5 fluorophore (E07-Cy5). The E07-Cy5 was incubated at a concentration of 100 nM for 15 minutes, 1 hour, 3 hours, and 24 hours, in the absence or presence of gefitinib (20 μM) on LN319 and U87 cell lines, following the same protocol used for cetuximab, except that nucleic acid aptamers were first denatured at 95 °C for 3 min, incubated on ice for 5 min before being resuspended at 100 nM final concentration.

Representative images in Figure 3A show the absence of E07-Cy5 aptamer in the EGFR-negative LN319 cell line as already shown (Cruz Da Silva et al., 2022). As for cetuximab, without treatment, E07-Cy5 is internalized in U87 cells, up to 3.4x more after 24h of treatment compared to 15 min. Upon gefitinib treatment, E07-Cy5 is further internalized, to 2.3x at 24h compared to untreated U87 cells at the same time point (Figure 3B). This gefitinib-mediated enhancement of endocytosis is associated with a morphological alteration of intracellular vesicles (Figure 3A). As for Cetuximab, we quantified the colocalization of the E07-Cy5 with early and late endosomes using Rab5 and CD63 immunolabeling. Our results show that the intracellular localization of the E07 aptamer in early and late endosomes remains essentially unchanged, irrespective of gefitinib treatment, proving that the anti-EGFR aptamer E07 is recruited to endosomes upon gefitinib treatment (Figure S3).

**Figure 3.**
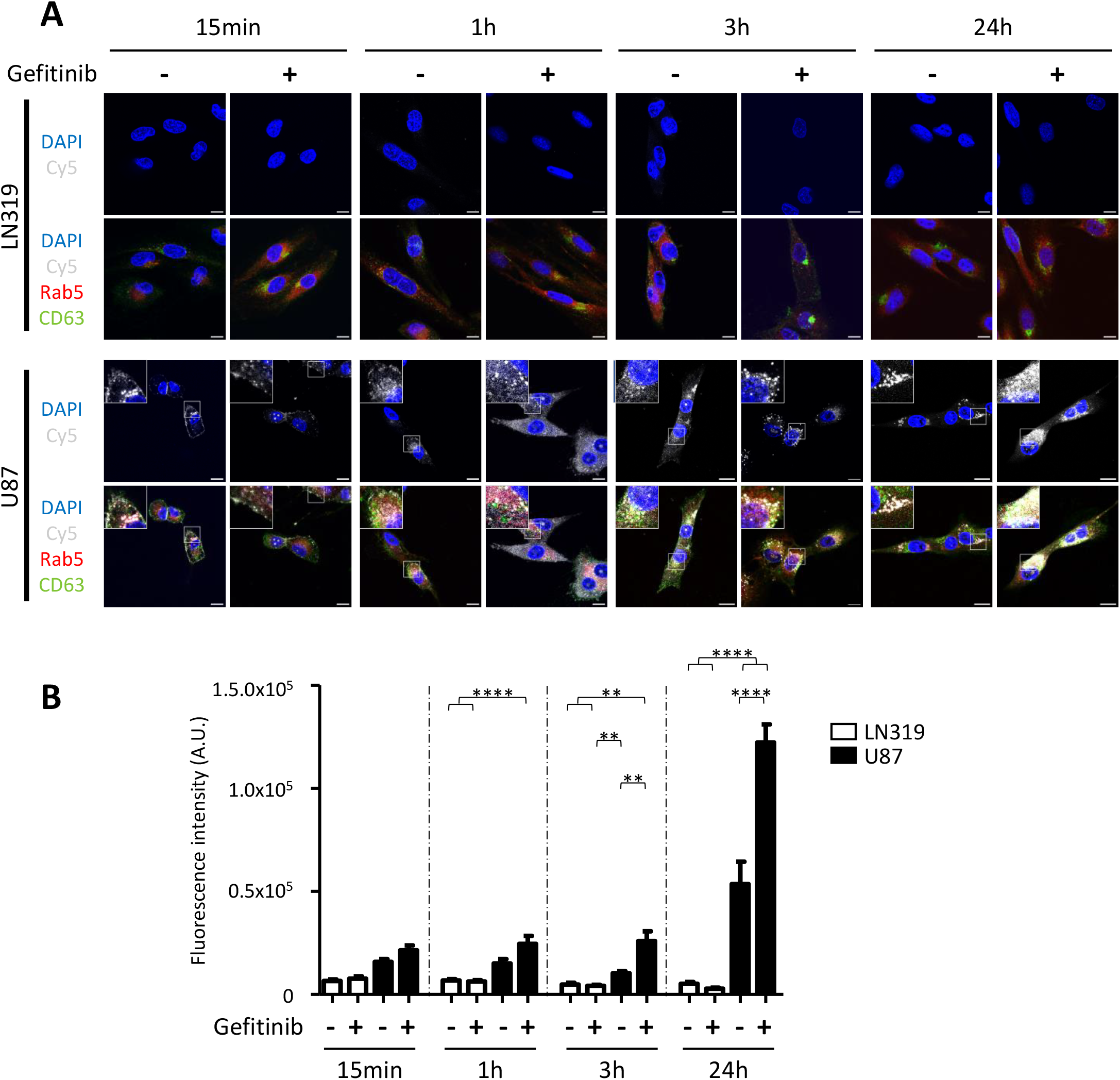
Effect of gefitinib treatment on the internalization and co-localisation of the cyanine5-conjugated aptamer E07 in GBM cell lines. The same experiment as for those described for cetuximab was used, except that the cyanine5-conjugated cetuximab was replaced with the cyanine5-conjugated aptamer E07 (E07-Cy5). **A**. Representative confocal images, showing DAPI (blue to label nucleus), Rab5 (red to label early endosomes), CD63 (green to label late endosomes), and cyanine5-conjugated aptamer E07 (grey). Magnified images of U87 cells are from the inserts. Scale bar = 12 μm. **B**. Quantification of the E07-Cy5 internalization. Fluorescence intensity is expressed in arbitrary units.

Bioimaging experiments realized with antibodies (Figure 2) and aptamers (Figure 3) both showed that their intracellular delivery is affected and disturbed from the very first hours of gefitinib treatment, and follow a very similar internalization profile and kinetic. For both ligands, gefitinib not only enhances antibodies and aptamers internalization shown by enhancement of fluorescence intensity inside the cells, but it is also remarkable that this accumulation results in larger endosomes compared to control cells. This modification in endosome morphology is a characteristic of GME, suggesting endosomes fusion and/or endosomal arrest, as we recently described both for EGFR and EGF internalization mediated by gefitinib (Blandin et al., 2021). But might this gefitinib-mediated enhancement of intracellular ligand delivery have a functional impact? To address this issue, we extended our investigations with functional studies to determine whether gefitinib could improve the toxicity of an ADC.

### 3.4. Gefitinib synergistically enhances antibody-drug conjugate efficacy and lowers dose requirements

We evaluated the pharmacological benefits of the TKI/ligand therapeutic combination, focusing on clinically used drugs and antibodies for treating diverse tumors, including GBM. The functional effect on cell viability of gefitinib was tested in association with an ADC composed of cetuximab-APN-PEG5-VC-MMAE, referred to as Cetuximab-MMAE below for simplification (with APN for arylpropiolonitrile, PEG for polyethylene glycol, VC for valine-citrulline, MMAE for monomethyl auristatin E), with drug-to-antibody-ratio of 4.3 and compared with cetuximab. Cell viability was quantified thanks to WST-1 assay (Houël et al., 2015). To confirm that the measured effect of treatments was cytototoxic and not cytostatic or linked to a decrease of metabolism, we followed the cultures by imaging using an automated imaging apparatus, the Incucyte-S3 (Sartorius). Images confirmed that the observed effects were due to a decrease of cell viability.

Figure 4A shows that gefitinib alone at 2 and 20 μM has no effect on U87 cell viability. In contrast, the highly cytotoxic drug MMAE, whether used alone at 1 μM or in combination with gefitinib, significantly affected cell viability. Similarly, chlorpromazine at 50 μM, used as an internal control of cytotoxicity (Calas et al., 2020; Dudani and Gupta, 1987), impacted strongly cell viability. These data confirm the consistency of the assay.

**Figure 4.**
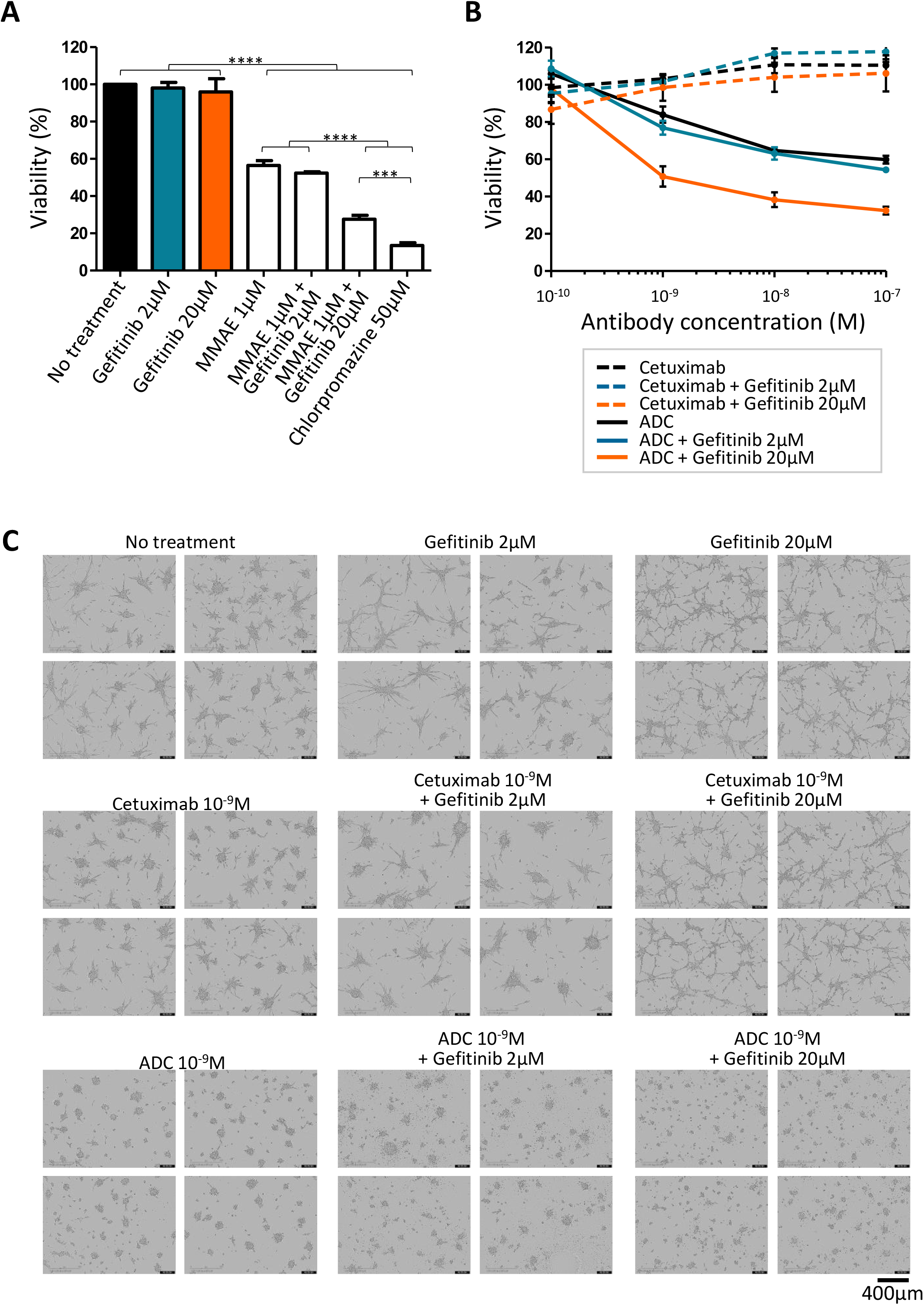
Synergistic activity of antibodies and gefitinib treatments on U87 cells viability. Cells were seeded for 24h, then treated for 72h, before quantifying their viability. **A**. Effect of control treatments on cell viability. Data for untreated cells are shown in black, gefitinib treatments at 2 and 20 µM are shown in blue and orange, respectively, and other treatments in white (MMAE 1 μM, MMAE 1 μM with gefitinib at 2 and 20 µM and chlorpromazine 50 μM). Data were reported as histograms, using Graphpad software. **B**. Effect on cell viability of cetuximab or ADC at different concentrations (10^−10^, 10^−9^, 10^−8^ and 10^−7^M) without gefitinib (black), with 2 μM gefitinib (blue) and with 20 μM gefitinib (orange). Cetuximab and ADC treatments are shown as dotted and plain lines, respectively. Statistical analysis is provided in Figure S4 and the different conditions with the 10^−9^ M antibody concentration are presented in bar graphs in Figure S5. **C**. Representative images from IncuCyte S3 of U87 cells at the end of the treatments with gefitinib 2 and 20 μM, cetuximab 10^−9^M ± gefitinib 2 and 20 μM, and ADC 10^−9^M ± gefitinib 2 and 20 μM. Four images / condition are shown. Scale = 400μm.

We then compared the effect of gefitinib treatment at 2 and 20 μM with Cetuximab or Cetuximab-MMAE, at concentrations ranging from 10^−10^ to 10^−7^ M. These results are shown in Figures 4B as curves and in Figure S4 as histograms. Our results with cetuximab show that irrespective of the concentration of cetuximab and the concomitant treatment with gefitinib, cell viability is barely significantly altered, indicating that under the conditions tested, cetuximab ± gefitinib treatment has no effect on U87 cell viability. The conjugation of MMAE to cetuximab allows the ADC to achieve cell death rates of 16 and 46% for 10^−9^ and 10^−7^ M of Cetuximab-MMAE, respectively. Addition of gefitinib at 2 μM to Cetuximab-MMAE does not modify the cell viability compared to Cetuximab-MMAE. However, it is particularly noteworthy that the addition of 20 μM Gefitinib significantly enhances the efficacy of Cetuximab-MMAE at concentrations ranging from 10^−9^-10^−7^ M, resulting in the death of 49-67% of the cells. At the concentration of 10^−9^ M, the addition of 20 μM Gefitinib further enhances cell death compared to using Cetuximab-MMAE alone at a concentration 100 times higher. Detailed results for an antibody concentration of 10^−9^ M are presented in Figure 4C (photographs) and Figure S5 (histograms). This indicates that a lower dose of the ADC combined with 20 μM Gefitinib results in higher cell death than significantly higher doses of the ADC alone.

## 4. DISCUSSION

EGFR is a key therapeutic target for GBM, but the targeted approaches have failed to improve patient care. The development of EGFR-targeted new ligands such as ADCs provides hope of therapeutic advances in the treatment of GBM. However, the development of these treatments has been slowed by serious secondary toxicity and considerable treatment costs. Gefitinib, an EGFR TKI used as anti-EGFR therapy in the treatment of lung cancer, does not benefit in GBM patients. In a previous study, we demonstrated in an *in vitro* GBM model that gefitinib and other TKIs directed against EGFR induce a massive internalization of EGFR, named gefitinib-mediated endocytosis (GME) (Blandin et al., 2021; Cruz Da Silva et al., 2021a). Here, in order to boost anti-tumor activities, we exploited this unexpected property of TKIs to propose a new approach using gefitinib to enhance the internalization of two types of ligands of therapeutic interest: antibodies and aptamers. Figure 5 summarizes the bioimaging and functional results from our study.

**Figure 5.**
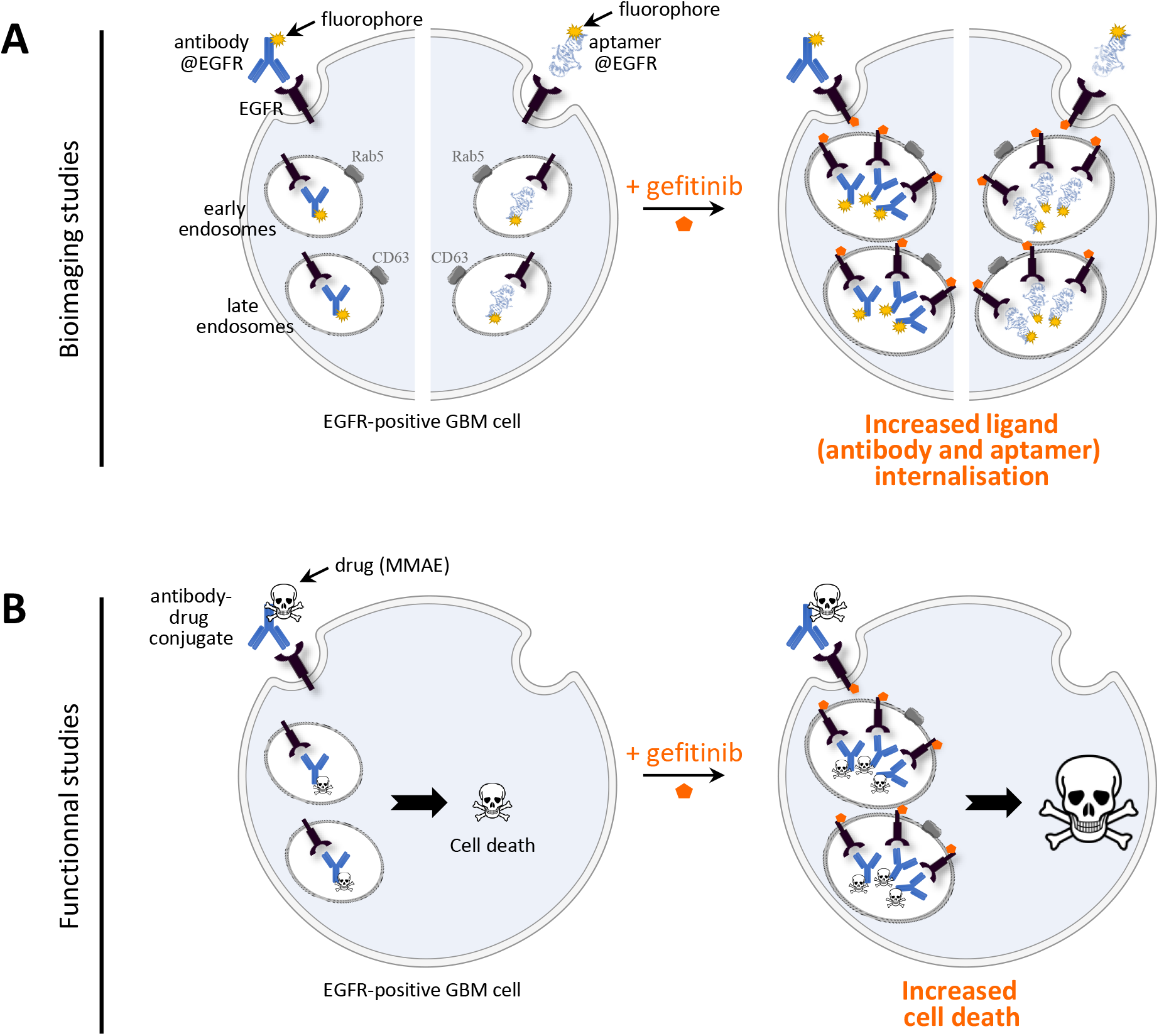
Schematic summarizing experiments and results obtained in this study, highlighting the effect of new therapeutic combination that enhance the internalization of ligands targeting EGFR in GBM cells. A. Scheme of bioimaging studies, illustrating the effect of gefitinib on the internalization of cetuximab and aptamer E07. Without gefitinib, the two fluorophore-conjugated EGFR-ligands, antibodies (shown to the left side of the cell) and aptamers (shown to the right side of the cell) are internalized in EGFR-positive GBM cells. Following gefitinib treatment, the internalization of the two ligands is further enhanced, symbolized by a larger quantity of antibodies and aptamers within the intracellular compartments. **B**. Functional studies realized with the antibody-drug conjugate cetuximab-MMAE, shows that gefitinib treatment enhances cell death compared to cells treated with the ADC alone.

Bioimaging with fluorescently labeled ligands revealed that GME enhances the rapid internalization of cetuximab and E07 aptamer, with longer gefitinib treatment leading to increased ligand internalization. After gefitinib treatment, ligands are localized in large endosomes, suggesting endosomes fusion and/or endosomal arrest (Blandin et al., 2021). A possible explanation is the involvement of gefitinib’s lysotropism in the mechanism of action of GME. Lysotropism refers to the tendency of certain compounds to accumulate within endolysosomes due to their physicochemical properties. Gefitinib exhibits lysotropic behavior leading to its accumulation in intracellular compartments where it can alter endolysosomal function. This accumulation may contribute to the drug’s efficacy by affecting cellular processes associated with endolysosomal activity (Hussein et al., 2021). Proteomic analysis showed that gefitinib treatment in lung carcinoma cell lines increased the expression or ubiquitination of proteins involved in endocytic pathways (Li et al., 2018). Neratinib and afatinib were also shown to increase polyubiquitination and subsequent internalization of the receptor HER2 in lung cancers (Li et al., 2020). GME could therefore be the consequence of a general disruption of membrane trafficking following accumulation of gefitinib in acidic endomembrane compartments.

In the present study, we did not conduct a mechanistical effect of GME. We rather focused on the possibility that the gefitinib-mediated prolonged accumulation of ligands in endosomes could promote endosomal escape thanks to endosome destabilization, an outcome of particular benefit for targeted therapy. Consequently, the bio-imaging studies were complemented with functional studies, applied on cetuximab and the ADC Cetuximab-MMAE. We showed that GME did not enhance the efficacy of cetuximab, but synergistically enhance the efficacy of the ADC. This effect is highly promising in therapeutic perspectives, as gefitinib treatment allows to reduce the ADC dose by 100-fold while preserving a synergistic efficacy on U87 cells. Minimizing dose is crucial to mitigate side effects, particularly since combined therapies, despite their enhanced cancer cell killing ability, often increase overall toxicity. This challenge is also significant for ADCs. Indeed, the failure of depatuxizumab mafodotin (ABT-414, depatux-m, AbbVie), an anti-EGFR ADC based of the monomethyl auristatinF (MMAF), tested in clinical phases II and III in GBM, highlights the major challenge of improving the efficacy of ADCs in their development, particularly in reducing the dose as MMAF-based therapies frequently induce ocular toxicities (Lassman et al., 2022). *In fine*, for GME as for other strategies, enhancing tumor cell sensitivity by improving ADC intracellular accumulation (Takahashi et al., 2022) should allow ADC dose reduction, thereby minimizing therapeutic side effects and significantly lowering costs. To our knowledge, no drug specifically targeting EGFR endocytosis has been approved for clinical use. Our observations may open the door to new clinical settings modulating EGFR trafficking.

The proposed approach combines two compounds, gefitinib and cetuximab, both of which are already in clinical use or in advanced clinical trials for many different tumors, and are generally safe to use concomitantly even with conventional radiation (Guimond et al., 2022). This could facilitate a faster transition to therapeutic applications. The combination of cetuximab/gefitinib, sometimes alongside other drugs, is not currently being tested in clinical trials for the treatment of GBM. However, it is being studied for other tumors, including Advanced/Metastatic Non-Small Cell Lung Cancer (NCT00162318), Colorectal Cancer / Head and Neck Cancer / Non-Small Cell Lung Cancer (NCT00820417), and Advanced or Recurrent Extrahepatic Cholangiocarcinoma and Gallbladder Carcinoma (NCT02836847, NCT03768375). Other strategies combining dual targeting of EGFR with monoclonal antibodies and TKIs are in clinical trials, as for example Cetuximab Plus Afatinib in Non-Small Cell Lung Cancer (NCT01090011), Refractory wtKRAS Metastatic Colorectal Cancer (NCT01919879), Advanced Solid Tumours (NCT02020577), or Cetuximab/Dasatinib (NCT00388427), Cetuximab/Erlotinib (NCT00895362), Cetuximab/ Regorafenib (NCT02095054) in Advanced malignancies. Such combinations might overcome drug resistance (Zhang et al., 2022). In GBM, clinical trials have explored or are currently investigating some EGFR-targeted therapies, such as tyrosine kinase inhibitors (gefitinib [NCT01310855 and NCT00250887]), antibodies (e.g. cetuximab [NCT00311857 and NCT00463073], panitumumab [NCT01017653] (Shikalov et al., 2024), antibody-drug conjugates (recently reviewed (High et al., 2024)). Clinical trials combining bevacizumab (an anti-VEGF antibody) and erlotinib have failed (NCT00525525, NCT00720356, NCT00671970) (Cruz Da Silva et al., 2021b). While cetuximab has not yet been used as a tumor-targeting antibody in anti-GBM ADCs, some of them in clinical trials (MRG003, HLX42) hold promise for targeting GBM (High et al., 2024). In preclinical settings, Li et al. recently reported that the cetuximab-MMAE AGCM-22 effectively inhibited GBM cell proliferation, and that both mouse and human orthotopic tumor models exhibited selective tumor-targeting efficacy with superior anti-GBM activity compared to temozolomide chemotherapy (Li et al., 2024). Our results reinforce the value of cetuximab as an ADC, highlighting the synergistic effect of combining cetuximab-based ADCs and gefitinib on EGFR-positive GBM. The two combined compounds target the same receptor, which could improve the overall selectivity of the co-treatment, after assessing EGFR expression in the tumor.

In recent years, scientists have become increasingly interested in using aptamers for drug delivery. Within this context, our study compared gefitinib-mediated antibody endocytosis with that of EGFR-targeting aptamers. Cetuximab and aptamer E07 were reported to exhibit the same binding affinity for EGFR in molecular binding experiments: 5.2 nM (Patel et al., 2007) and 2.4 nM (Li et al., 2011), respectively, and in our bioimaging experiments, they were used at comparable concentrations: 66 nM for cetuximab and 100 nM for aptamer E07. Under these conditions, we demonstrated that both ligands exhibit a very similar internalization profile, both with and without gefitinib treatment. Gefitinib is therefore likely to be of major interest in promoting aptamer internalization when used as a targeting ligand, following the approach of antibody conjugates.

## 5. CONCLUSION

Gefitinib not only enhanced the internalization of two specific EGFR ligands, cetuximab and aptamer E07, but also has a synergistic effect on the cytotoxic efficacy of Cetuximab-MMAE ADC in EGFR-expressing GBM cells. This study highlights the importance of developing combined therapy, and is part of a broader project aimed at improving antibodies and aptamers endocytosis and consequently their therapeutic efficacy in human clinics. Future validation of the GME/ADC combination *ex vivo* in tumoroids and *in vivo* in murine tumour xenograft models, along with experiments to confirm the potential of GME for aptamer-conjugates might support a new treatment strategy for GBM, and other cancers where anti-EGFR therapies are considered. Given that this strategy is based on some compounds already in clinical use or in clinical trials, it could be rapidly translated into clinical practice.

## Abbreviations

ADC: antibody-drug conjugate
EGFR: epidermal growth factor receptor
GBM: glioblastoma multiform
GME: gefitinib-mediated endocytosis
mAb: monoclonal antibody
MMAE: monométhyl auristatine E
TKI: tyrosine kinase inhibitor.

## AUTHOR CONTRIBUTIONS

Conceptualization, E.C.D.S., L.C., M.L.; methodology, E.C.D.S., H.J., C.D.A., D.S., V.C., P.V., M.L. and L.C.; formal analysis, E.C.D.S., H.J., C.D.A., D.S., R.V., V.C., P.V., M.L. and L.C.; investigation, E.C.D.S., M.L. and L.C.; writing—original draft preparation, E.C.D.S. and L.C.; writing, L.C.; project administration, M.L. and L.C.; funding acquisition, M.L. and L.C. All authors have read and agreed to this version of the manuscript.

## DATA AVAILABILITY STATEMENT

Data supporting reported results will be provided by corresponding author upon request.

## REFERENCES

Blandin, A.-F., Cruz Da Silva, E., Mercier, M.-C., Glushonkov, O., Didier, P., Dedieu, S., Schneider, C., Devy, J., Etienne-Selloum, N., Dontenwill, M., Choulier, L., Lehmann, M., 2021. Gefitinib induces EGFR and α5β1 integrin co-endocytosis in glioblastoma cells. Cell. Mol. Life Sci. 78, 2949–2962. 10.1007/s00018-020-03686-6

Bolte, S., Cordelières, F.P., 2006. A guided tour into subcellular colocalization analysis in light microscopy. Journal of Microscopy 224, 213–232. 10.1111/j.1365-2818.2006.01706.x

Calas, A.-G., Hanak, A.-S., Jaffré, N., Nervo, A., Dias, J., Rousseau, C., Courageux, C., Brazzolotto, X., Villa, P., Obrecht, A., Goossens, J.-F., Landry, C., Hachani, J., Gosselet, F., Dehouck, M.-P., Yerri, J., Kliachyna, M., Baati, R., Nachon, F., 2020. Efficacy Assessment of an Uncharged Reactivator of NOP-Inhibited Acetylcholinesterase Based on Tetrahydroacridine Pyridine-Aldoxime Hybrid in Mouse Compared to Pralidoxime. Biomolecules 10, 858. 10.3390/biom10060858

Caldieri, G., Malabarba, M.G., Di Fiore, P.P., Sigismund, S., 2018. EGFR Trafficking in Physiology and Cancer. Prog Mol Subcell Biol 57, 235–272. 10.1007/978-3-319-96704-2_9

Chang, X., Izumchenko, E., Solis, L.M., Kim, M.S., Chatterjee, A., Ling, S., Monitto, C.L., Harari, P.M., Hidalgo, M., Goodman, S.N., Wistuba, I.I., Bedi, A., Sidransky, D., 2013. The Relative Expression of Mig6 and EGFR Is Associated with Resistance to EGFR Kinase Inhibitors. PLOS ONE 8, e68966. 10.1371/journal.pone.0068966

Colin, M., Delporte, C., Janky, R., Lechon, A.-S., Renard, G., Van Antwerpen, P., Maltese, W.A., Mathieu, V., 2019. Dysregulation of Macropinocytosis Processes in Glioblastomas May Be Exploited to Increase Intracellular Anti-Cancer Drug Levels: The Example of Temozolomide. Cancers 11, 411. 10.3390/cancers11030411

Cruz Da Silva, E., Choulier, L., Thevenard-Devy, J., Schneider, C., Carl, P., Rondé, P., Dedieu, S., Lehmann, M., 2021a. Role of Endocytosis Proteins in Gefitinib-Mediated EGFR Internalisation in Glioma Cells. Cells 10, 3258. 10.3390/cells10113258

Cruz Da Silva, E., Foppolo, S., Lhermitte, B., Ingremeau, M., Justiniano, H., Klein, L., Chenard, M.-P., Vauchelles, R., Abdallah, B., Lehmann, M., Etienne-Selloum, N., Dontenwill, M., Choulier, L., 2022. Bioimaging Nucleic-Acid Aptamers with Different Specificities in Human Glioblastoma Tissues Highlights Tumoral Heterogeneity. Pharmaceutics 14, 1980. 10.3390/pharmaceutics14101980

Cruz Da Silva, E., Mercier, M.-C., Etienne-Selloum, N., Dontenwill, M., Choulier, L., 2021b. A Systematic Review of Glioblastoma-Targeted Therapies in Phases II, III, IV Clinical Trials. Cancers (Basel) 13. 10.3390/cancers13081795

DeVay, R.M., Delaria, K., Zhu, G., Holz, C., Foletti, D., Sutton, J., Bolton, G., Dushin, R., Bee, C., Pons, J., Rajpal, A., Liang, H., Shelton, D., Liu, S.-H., Strop, P., 2017. Improved Lysosomal Trafficking Can Modulate the Potency of Antibody Drug Conjugates. Bioconjugate Chem. 28, 1102–1114. 10.1021/acs.bioconjchem.7b00013

Diamantis, N., Banerji, U., 2016. Antibody-drug conjugates--an emerging class of cancer treatment. Br J Cancer 114, 362–367. 10.1038/bjc.2015.435

Dudani, A.K., Gupta, R.S., 1987. Effect of chlorpromazine and trifluoperazine on cytoskeletal components and mitochondria in cultured mammalian cells. Tissue Cell 19, 183–196. 10.1016/0040-8166(87)90004-8

Dunn, K.W., Kamocka, M.M., McDonald, J.H., 2011. A practical guide to evaluating colocalization in biological microscopy. Am J Physiol Cell Physiol 300, C723–C742. 10.1152/ajpcell.00462.2010

Fares, J., Wan, Y., Mair, R., Price, S.J., 2024. Molecular diversity in isocitrate dehydrogenase-wild-type glioblastoma. Brain Commun 6, fcae108. 10.1093/braincomms/fcae108

Fechter, P., Cruz Da Silva, E., Mercier, M.-C., Noulet, F., Etienne-Seloum, N., Guenot, D., Lehmann, M., Vauchelles, R., Martin, S., Lelong-Rebel, I., Ray, A.-M., Seguin, C., Dontenwill, M., Choulier, L., 2019. RNA Aptamers Targeting Integrin α5β1 as Probes for Cyto- and Histofluorescence in Glioblastoma. Mol Ther Nucleic Acids 17, 63–77. 10.1016/j.omtn.2019.05.006

Furnari, F.B., Cloughesy, T.F., Cavenee, W.K., Mischel, P.S., 2015. Heterogeneity of epidermal growth factor receptor signalling networks in glioblastoma. Nat Rev Cancer 15, 302–310. 10.1038/nrc3918

Gan, Hui K., Parakh, S., Osellame, L.D., Cher, L., Uccellini, A., Hafeez, U., Menon, S., Scott, A.M., 2023. Antibody drug conjugates for glioblastoma: current progress towards clinical use. Expert Opinion on Biological Therapy.

Gan, Hui K, Parakh, S., Osellame, L.D., Cher, L., Uccellini, A., Hafeez, U., Menon, S., Scott, A.M., 2023. Antibody drug conjugates for glioblastoma: current progress towards clinical use. Expert Opinion on Biological Therapy 23, 1089–1102. 10.1080/14712598.2023.2282729

Grandal, M.V., Zandi, R., Pedersen, M.W., Willumsen, B.M., van Deurs, B., Poulsen, H.S., 2007. EGFRvIII escapes down-regulation due to impaired internalization and sorting to lysosomes. Carcinogenesis 28, 1408–1417. 10.1093/carcin/bgm058

Guimond, E., Tsai, C.J., Hosni, A., O’Kane, G., Yang, J., Barry, A., 2022. Safety and Tolerability of Metastasis-Directed Radiation Therapy in the Era of Evolving Systemic, Immune, and Targeted Therapies. Adv Radiat Oncol 7, 101022. 10.1016/j.adro.2022.101022

High, P., Guernsey, C., Subramanian, S., Jacob, J., Carmon, K.S., 2024. The Evolving Paradigm of Antibody–Drug Conjugates Targeting the ErbB/HER Family of Receptor Tyrosine Kinases. Pharmaceutics 16, 890. 10.3390/pharmaceutics16070890

Houël, E., Fleury, M., Odonne, G., Nardella, F., Bourdy, G., Vonthron-Sénécheau, C., Villa, P., Obrecht, A., Eparvier, V., Deharo, E., Stien, D., 2015. Antiplasmodial and anti-inflammatory effects of an antimalarial remedy from the Wayana Amerindians, French Guiana: takamalaimë (Psidium acutangulum Mart. ex DC., Myrtaceae). J Ethnopharmacol 166, 279–285. 10.1016/j.jep.2015.03.015

Hussein, N.A., Malla, S., Pasternak, M.A., Terrero, D., Brown, N.G., Ashby, C.R., Assaraf, Y.G., Chen, Z.-S., Tiwari, A.K., 2021. The role of endolysosomal trafficking in anticancer drug resistance. Drug Resist Updat 57, 100769. 10.1016/j.drup.2021.100769

Inda, M.-M., Bonavia, R., Mukasa, A., Narita, Y., Sah, D.W.Y., Vandenberg, S., Brennan, C., Johns, T.G., Bachoo, R., Hadwiger, P., Tan, P., DePinho, R.A., Cavenee, W., Furnari, F., 2010. Tumor heterogeneity is an active process maintained by a mutant EGFR-induced cytokine circuit in glioblastoma. Genes Dev. 24, 1731–1745. 10.1101/gad.1890510

Kalim, M., Chen, J., Wang, S., Lin, C., Ullah, S., Liang, K., Ding, Q., Chen, S., Zhan, J., 2017. Intracellular trafficking of new anticancer therapeutics: antibody–drug conjugates. Drug Des Devel Ther 11, 2265–2276. 10.2147/DDDT.S135571

Kondapalli, K.C., Llongueras, J.P., Capilla-González, V., Prasad, H., Hack, A., Smith, C., Guerrero-Cázares, H., Quiñones-Hinojosa, A., Rao, R., 2015. A Leak Pathway for Luminal Protons in Endosomes Drives Oncogenic Signaling in Glioblastoma. Nat Commun 6, 6289. 10.1038/ncomms7289

Kratschmer, C., Levy, M., 2018. Targeted Delivery of Auristatin-Modified Toxins to Pancreatic Cancer Using Aptamers. Molecular Therapy - Nucleic Acids 10, 227–236. 10.1016/j.omtn.2017.11.013

Lassman, A.B., Pugh, S.L., Wang, T.J.C., Aldape, K., Gan, H.K., Preusser, M., Vogelbaum, M.A., Sulman, E.P., Won, M., Zhang, P., Moazami, G., Macsai, M.S., Gilbert, M.R., Bain, E.E., Blot, V., Ansell, P.J., Samanta, S., Kundu, M.G., Armstrong, T.S., Wefel, J.S., Seidel, C., de Vos, F.Y., Hsu, S., Cardona, A.F., Lombardi, G., Bentsion, D., Peterson, R.A., Gedye, C., Bourg, V., Wick, A., Curran, W.J., Mehta, M.P., 2022. Depatuxizumab mafodotin in EGFR-amplified newly diagnosed glioblastoma: A phase III randomized clinical trial. Neuro Oncol 25, 339–350. 10.1093/neuonc/noac173

Li, B.T., Michelini, F., Misale, S., Cocco, E., Baldino, L., Cai, Y., Shifman, S., Tu, H.-Y., Myers, M.L., Xu, C., Mattar, M., Khodos, I., Little, M., Qeriqi, B., Weitsman, G., Wilhem, C.J., Lalani, A.S., Diala, I., Freedman, R.A., Lin, N.U., Solit, D.B., Berger, M.F., Barber, P.R., Ng, T., Offin, M., Isbell, J.M., Jones, D.R., Yu, H.A., Thyparambil, S., Liao, W.-L., Bhalkikar, A., Cecchi, F., Hyman, D.M., Lewis, J.S., Buonocore, D.J., Ho, A.L., Makker, V., Reis-Filho, J.S., Razavi, P., Arcila, M.E., Kris, M.G., Poirier, J.T., Shen, R., Tsurutani, J., Ulaner, G.A., de Stanchina, E., Rosen, N., Rudin, C.M., Scaltriti, M., 2020. HER2-Mediated Internalization of Cytotoxic Agents in ERBB2 Amplified or Mutant Lung Cancers. Cancer Discov 10, 674–687. 10.1158/2159-8290.CD-20-0215

Li, D., Sun, X., Li, Y., Shang, C., Dong, Y., Zhao, R., Zhang, H., Wang, Z., Fan, S., Ma, C., Li, X., 2024. AGCM-22, a novel cetuximab-based EGFR-targeting antibody-drug-conjugate with highly selective anti-glioblastoma efficacy. Bioorg Med Chem 102, 117657. 10.1016/j.bmc.2024.117657

Li, N., Nguyen, H.H., Byrom, M., Ellington, A.D., 2011. Inhibition of Cell Proliferation by an Anti-EGFR Aptamer. PLoS One 6. 10.1371/journal.pone.0020299

Li, W., Wang, H., Yang, Y., Zhao, T., Zhang, Z., Tian, Y., Shi, Z., Peng, X., Li, F., Feng, Y., Zhang, L., Jiang, G., Zhang, F., 2018. Integrative Analysis of Proteome and Ubiquitylome Reveals Unique Features of Lysosomal and Endocytic Pathways in Gefitinib-Resistant Non-Small Cell Lung Cancer Cells. PROTEOMICS 18, 1700388. 10.1002/pmic.201700388

Liu, Y., Li, Z., Zhang, M., Zhou, H., Wu, X., Zhong, J., Xiao, F., Huang, N., Yang, X., Zeng, R., Yang, L., Xia, Z., Zhang, N., 2021. Rolling-translated EGFR variants sustain EGFR signaling and promote glioblastoma tumorigenicity. Neuro Oncol 23, 743–756. 10.1093/neuonc/noaa279

Mahmoudian, F., Ahmari, A., Shabani, S., Sadeghi, B., Fahimirad, S., Fattahi, F., 2024. Aptamers as an approach to targeted cancer therapy. Cancer Cell International 24, 108. 10.1186/s12935-024-03295-4

Menon, S., Parakh, S., Scott, A.M., Gan, H.K., 2022. Antibody-drug conjugates: beyond current approvals and potential future strategies. Explor Target Antitumor Ther. 3, 252–277. 10.37349/etat.2022.00082

Mercier, M.-C., Dontenwill, M., Choulier, L., 2017. Selection of Nucleic Acid Aptamers Targeting Tumor Cell-Surface Protein Biomarkers. Cancers (Basel) 9. 10.3390/cancers9060069

Mullard, A., 2023. FDA approves second RNA aptamer. Nature Reviews Drug Discovery 22, 774–774. 10.1038/d41573-023-00148-z

Nishikawa, R., Ji, X.D., Harmon, R.C., Lazar, C.S., Gill, G.N., Cavenee, W.K., Huang, H.J., 1994. A mutant epidermal growth factor receptor common in human glioma confers enhanced tumorigenicity. Proc. Natl. Acad. Sci. U.S.A. 91, 7727–7731.

Pan, P.C., Magge, R.S., 2020. Mechanisms of EGFR Resistance in Glioblastoma. Int J Mol Sci 21, 8471. 10.3390/ijms21228471

Park, J.-W., Wollmann, G., Urbiola, C., Fogli, B., Florio, T., Geley, S., Klimaschewski, L., 2018. Sprouty2 enhances the tumorigenic potential of glioblastoma cells. Neuro Oncol 20, 1044–1054. 10.1093/neuonc/noy028

Patel, D., Lahiji, A., Patel, S., Franklin, M., Jimenez, X., Hicklin, D.J., Kang, X., 2007. Monoclonal Antibody Cetuximab Binds to and Down-regulates Constitutively Activated Epidermal Growth Factor Receptor vIII on the Cell Surface. Anticancer Research 27, 3355–3366.

Ray, P., Cheek, M.A., Sharaf, M.L., Li, N., Ellington, A.D., Sullenger, B.A., Shaw, B.R., White, R.R., 2012. Aptamer-Mediated Delivery of Chemotherapy to Pancreatic Cancer Cells. Nucleic Acid Ther 22, 295–305. 10.1089/nat.2012.0353

Shikalov, A., Koman, I., Kogan, N.M., 2024. Targeted Glioma Therapy—Clinical Trials and Future Directions. Pharmaceutics 16, 100. 10.3390/pharmaceutics16010100

Singh, S., Sadhukhan, S., Sonawane, A., 2023. 20 years since the approval of first EGFR-TKI, gefitinib: Insight and foresight. Biochim Biophys Acta Rev Cancer 1878, 188967. 10.1016/j.bbcan.2023.188967

Sousa, L.P., Lax, I., Shen, H., Ferguson, S.M., Camilli, P.D., Schlessinger, J., 2012. Suppression of EGFR endocytosis by dynamin depletion reveals that EGFR signaling occurs primarily at the plasma membrane. Proc Natl Acad Sci U S A 109, 4419–4424. 10.1073/pnas.1200164109

Stupp, R., Mason, W.P., van den Bent, M.J., Weller, M., Fisher, B., Taphoorn, M.J.B., Belanger, K., Brandes, A.A., Marosi, C., Bogdahn, U., Curschmann, J., Janzer, R.C., Ludwin, S.K., Gorlia, T., Allgeier, A., Lacombe, D., Cairncross, J.G., Eisenhauer, E., Mirimanoff, R.O., 2005. Radiotherapy plus Concomitant and Adjuvant Temozolomide for Glioblastoma. New England Journal of Medicine 352, 987–996. 10.1056/NEJMoa043330

Takahashi, J., Nakamura, S., Onuma, I., Zhou, Y., Yokoyama, S., Sakurai, H., 2022. Synchronous intracellular delivery of EGFR-targeted antibody–drug conjugates by p38-mediated non-canonical endocytosis. Sci Rep 12, 11561. 10.1038/s41598-022-15838-8

Walsh, A.M., Kapoor, G.S., Buonato, J.M., Mathew, L.K., Bi, Y., Davuluri, R.V., Martinez-Lage, M., Simon, M.C., O’Rourke, D.M., Lazzara, M.J., 2015. Sprouty2 Drives Drug Resistance and Proliferation in Glioblastoma. Molecular Cancer Research 13, 1227–1237. 10.1158/1541-7786.MCR-14-0183-T

Walsh, A.M., Lazzara, M.J., 2013. Regulation of EGFR trafficking and cell signaling by Sprouty2 and MIG6 in lung cancer cells. Journal of Cell Science 126, 4339–4348. 10.1242/jcs.123208

Westphal, M., Maire, C.L., Lamszus, K., 2017. EGFR as a Target for Glioblastoma Treatment: An Unfulfilled Promise. CNS Drugs 31, 723–735. 10.1007/s40263-017-0456-6

Xiang, D., Zheng, C., Zhou, S.-F., Qiao, S., Tran, P.H.-L., Pu, C., Li, Y., Kong, L., Kouzani, A.Z., Lin, J., Liu, K., Li, L., Shigdar, S., Duan, W., 2015. Superior Performance of Aptamer in Tumor Penetration over Antibody: Implication of Aptamer-Based Theranostics in Solid Tumors. Theranostics 5, 1083–1097. 10.7150/thno.11711

Yang, W., Wu, P.-F., Ma, J.-X., Liao, M.-J., Wang, X.-H., Xu, L.-S., Xu, M.-H., Yi, L., 2019. Sortilin promotes glioblastoma invasion and mesenchymal transition through GSK-3β/β-catenin/twist pathway. Cell Death Dis 10, 208. 10.1038/s41419-019-1449-9

Ying, H., Zheng, H., Scott, K., Wiedemeyer, R., Yan, H., Lim, C., Huang, J., Dhakal, S., Ivanova, E., Xiao, Y., Zhang, H., Hu, J., Stommel, J.M., Lee, M.A., Chen, A.-J., Paik, J.-H., Segatto, O., Brennan, C., Elferink, L.A., Wang, Y.A., Chin, L., DePinho, R.A., 2010. Mig-6 controls EGFR trafficking and suppresses gliomagenesis. Proc Natl Acad Sci U S A 107, 6912–6917. 10.1073/pnas.0914930107

Zhang, G., Yan, B., Guo, Y., Yang, H., Li, X., Li, J., 2022. Case Report: A patient with the rare third-generation TKI-resistant mutation EGFR L718Q who responded to afatinib plus cetuximab combination therapy. Front Oncol 12, 995624. 10.3389/fonc.2022.995624

Zhang, J.H., Chung, T.D., Oldenburg, K.R., 1999. A Simple Statistical Parameter for Use in Evaluation and Validation of High Throughput Screening Assays. J Biomol Screen 4, 67–73. 10.1177/108705719900400206

Zhou, G., Wilson, G., Hebbard, L., Duan, W., Liddle, C., George, J., Qiao, L., 2016. Aptamers: A promising chemical antibody for cancer therapy. Oncotarget 7, 13446–13463. 10.18632/oncotarget.7178

Zhou, X., Xie, S., Wu, S., Qi, Y., Wang, Z., Zhang, H., Lu, D., Wang, X., Dong, Y., Liu, G., Yang, D., Shi, Q., Bian, W., Yu, R., 2017. Golgi phosphoprotein 3 promotes glioma progression via inhibiting Rab5-mediated endocytosis and degradation of epidermal growth factor receptor. Neuro Oncol 19, 1628–1639. 10.1093/neuonc/nox104

