## Supplemental information for "New therapeutic combination to enhance endocytosis of antibodies and nucleic-acid aptamers targeting EGFR in glioblastoma cells"

### SUPPLEMENTAL LEGENDS

**Figure S1. Immunoblot showing EGFR expression at different incubation times with gefitinib in U87, T98 and LN443 cell lines.** Cells were treated for different times (0h, 4h and 24h) with 20 µM gefitinib. GAPDH was used as the loading control.

**Figure S2. Quantification of cetuximab/Rab5 and cetuximab/CD63 co-localization with or without gefitinib treatment**. The graphs represent the quantification of colocalized pixels using a using JACoP plugin ImageJ software (Bolte and Cordelières, 2006; Dunn et al., 2011) and show the colocalization of Cetuximab-Cy5 with Rab5 (left) and with CD63 (right). Data reported as Tukey box and whiskers were determined with the Pearson's correlation coefficient on LN319 (in white) and on U87 (in black) cells.

**Figure S3. Quantification of E07/Rab5 and E07/CD63 co-localisation with or without gefitinib treatment.** The graphs represent the quantification of colocalized pixels using a using JACoP plugin ImageJ software (Bolte and Cordelières, 2006; Dunn et al., 2011) and show the colocalization of E07-Cy5 with Rab5 (left) and with CD63 (right). Data reported as Tukey boxes were determined with the Pearson's correlation coefficient on LN319 (in white) and on U87 (in black) cells.

**Figure S4. Synergistic activity of antibodies and gefitinib treatments on U87 cells viability**. These data are identical to those shown in Figure 4, except that there are represented as histograms with statistics. Cetuximab and ADC treatments are shown as stripped and plain bars, respectively. Data for untreated cells are shown in black, gefitinib treatments at 2 and 20 µM are shown in blue and orange, respectively.

**Figure S5. Synergistic activity of antibodies at the concentration of $10^{-9}$ M and gefitinib treatments on U87 cells viability**. These data are identical to those shown in Figure 4 and S4, except that there are represented as histograms with statistics, only for the $10^{-9}$M antibody concentration. Cetuximab and ADC treatments are shown as stripped and plain bars, respectively. Data for untreated cells are shown in black, gefitinib treatments at 2 and 20 µM are shown in blue and orange, respectively. * $p \leq 0.05$, ** $p \leq 0.01$, *** $p \leq 0.001$, **** $p \leq 0.0001$.

### SUPPLEMENTAL METHODS

#### Western blot

Cells were lysed in 1% Triton X-100, NaF [100 mmol/L], NaPPi [10 mmol/L], and $Na_3VO_4$ [1 mmol/L] in PBS, supplemented with complete anti-protease cocktail (Roche, Basel, Switzerland). A total of 10 µg of protein was separated on precast gradient 4–20% SDS-PAGE gels (Bio-Rad, Hercules, CA, USA) and transferred to polyvinylidene fluoride (PVDF) membranes (Amersham Bioscience, Buckinghamshire, UK). After blocking, membranes were probed with primary antibodies targeting EGFR (D38B1, #4267; Cell Signaling Technology, Danvers, MA, USA), and glyceraldehyde 3-phosphate dehydrogenase (GAPDH, clone 6C5, Millipore, Molsheim, France). Immunological complexes were revealed with horseradish peroxidase (HRP)-conjugated secondary antibodies (Promega, Madison, WI, USA) at a 1/10,000 dilution. Revelation was performed with enhanced chemiluminescence (ECL; BioRad) using an LAS4000 imager (GE Healthcare, Dornstadt, Germany). GAPDH was used as housekeeping protein to serve as the loading control for all cell lysate samples. The quantification of non-saturated images was performed with ImageJ software.

### SUPPLEMENTAL REFERENCES

Bolte, S., Cordelières, F.P., 2006. A guided tour into subcellular colocalization analysis in light microscopy. Journal of Microscopy 224, 213–232. https://doi.org/10.1111/j.1365-2818.2006.01706.x

Dunn, K.W., Kamocka, M.M., McDonald, J.H., 2011. A practical guide to evaluating colocalization in biological microscopy. Am J Physiol Cell Physiol 300, C723–C742. https://doi.org/10.1152/ajpcell.00462.2010

### Figure S1

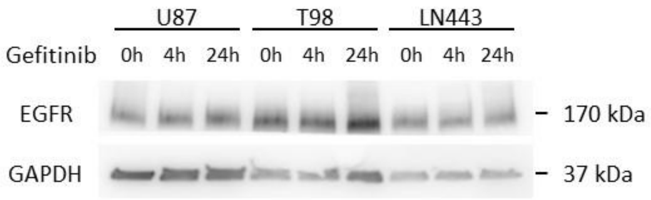


### Figure S2

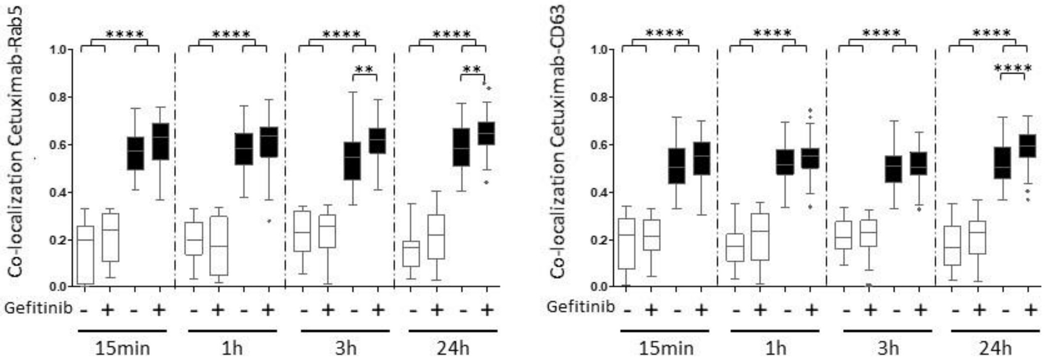


### Figure S3

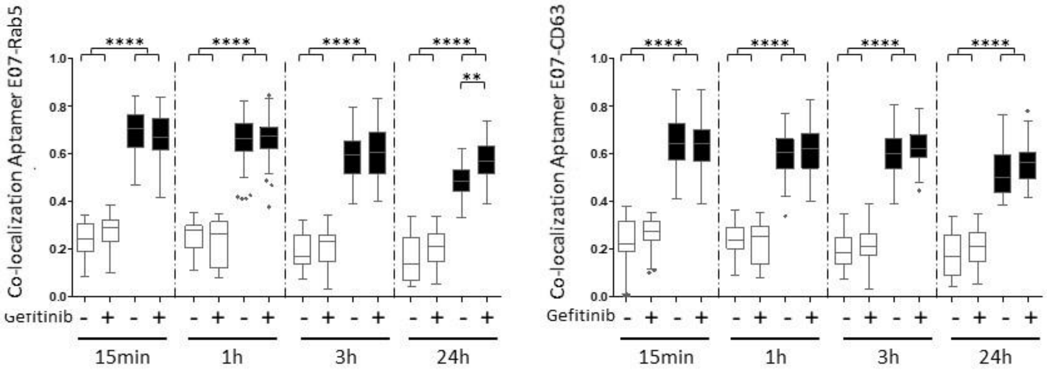

Figure S4

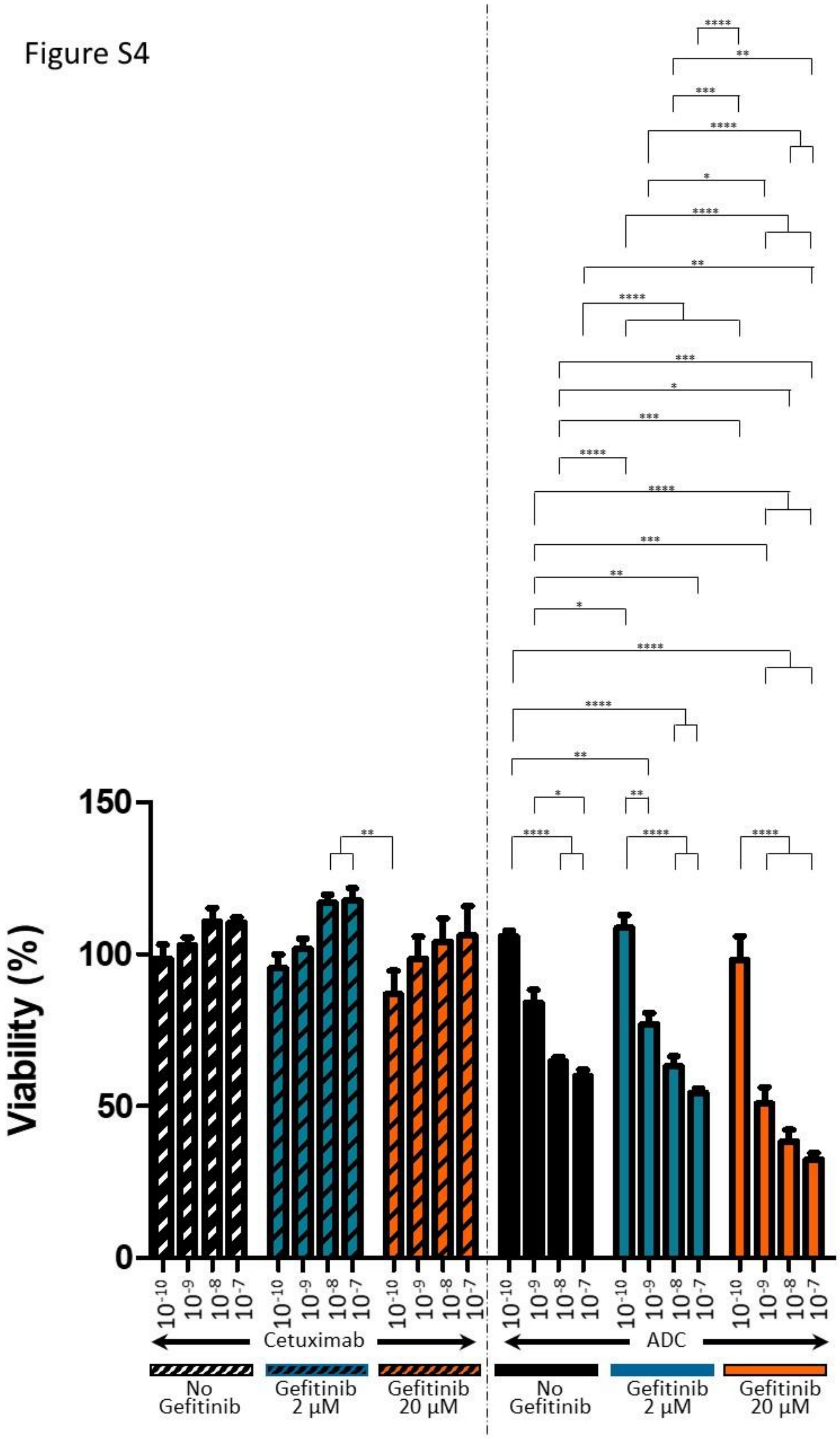

Viability (%)
150
100
50
0
10-10 10-9 10-8 10-7
Cetuximab
ADC
No Gefitinib
Gefitinib 2 µM
Gefitinib 20 µM

Figure S5

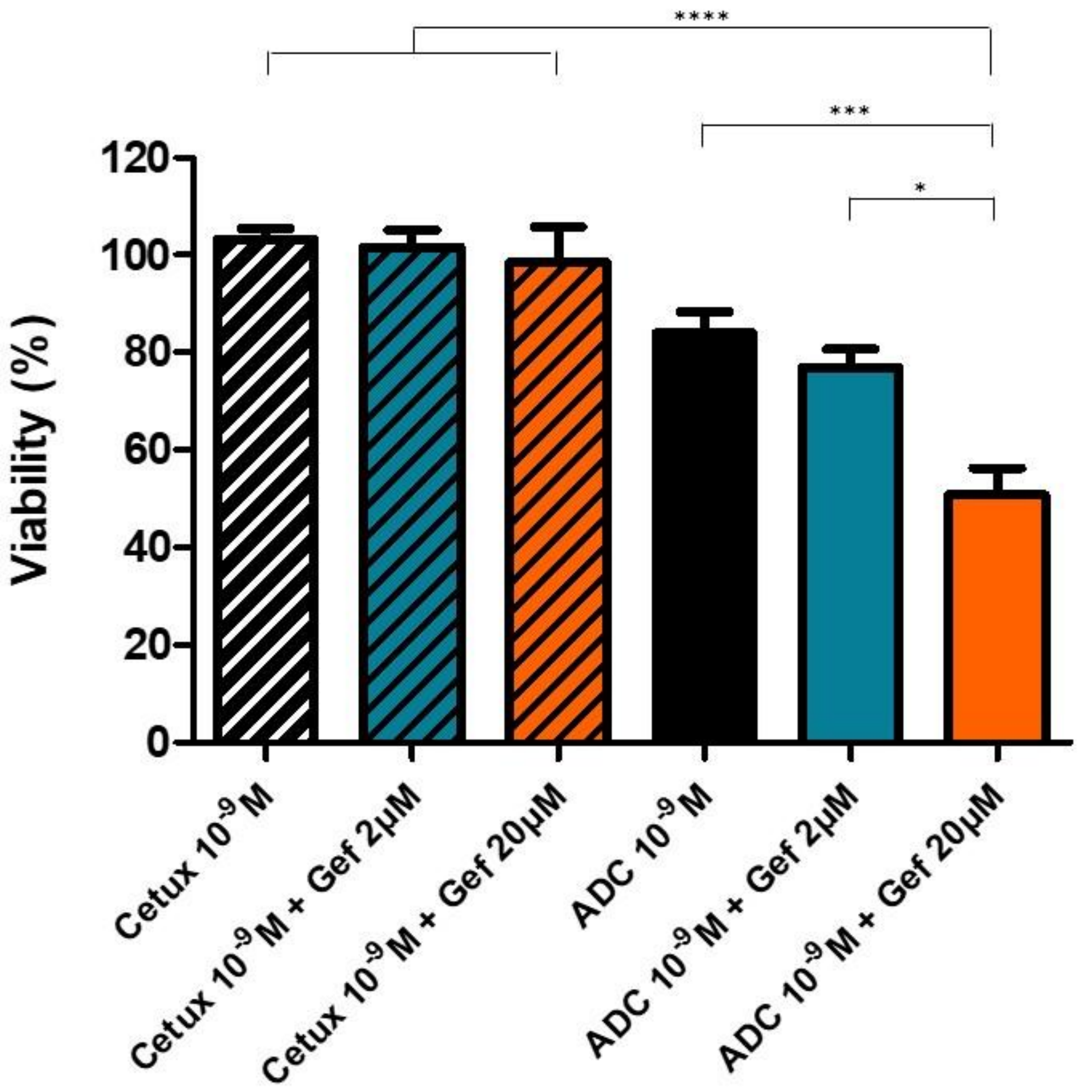
****
***
*
120
100
80
60
40
20
0
Viability (%)
Cetux 10-9 M
Cetux 10-9 M + Gef 2µM
Cetux 10-9 M + Gef 20µM
ADC 10-9 M
ADC 10-9 M + Gef 2µM
ADC 10-9 M + Gef 20µM
